# Graft incompatibility between pepper and tomato can be attributed to genetic incompatibility between diverged immune systems

**DOI:** 10.1101/2024.03.29.587379

**Authors:** Hannah Rae Thomas, Alice Gevorgyan, Alexandra Hermanson, Samantha Yanders, Lindsay Erndwein, Matthew Norman-Ariztía, Erin E. Sparks, Margaret H Frank

## Abstract

- Graft compatibility is the capacity of two plants to form cohesive vascular connections. Tomato and pepper are incompatible graft partners; however, the underlying cause of graft rejection between these two species remains unknown.
- We diagnosed graft incompatibility between tomato and diverse pepper varieties based on weakened biophysical stability, decreased growth, and persistent cell death using trypan blue and TUNEL assays. Transcriptomic analysis of cell death in the junction was performed using RNA-sequencing, and molecular signatures for incompatible graft response were characterized based on meta-transcriptomic comparisons with other biotic processes.
- We show that tomato is broadly incompatible with diverse pepper cultivars. These incompatible graft partners activate prolonged transcriptional changes that are highly enriched for defense processes. Amongst these processes was broad NLR upregulation and hypersensitive response. Using transcriptomic datasets for a variety of biotic stress treatments, we identified a significant overlap in the genetic profile of incompatible grafting and plant parasitism. In addition, we found over 1000 genes that are uniquely upregulated in incompatible grafts.
- Based on NLR overactivity, DNA damage, and prolonged cell death we have determined that tomato and pepper graft incompatibility is likely caused by a form of genetic incompatibility, which triggers a hyperimmune-response.

## Introduction

Grafting is an ancient agricultural practice that is used to propagate plants and combine desirable traits between independent root and shoot systems (Harrison & Burgess, 1962; Mudge *et al*., 2009; Warschefsky *et al*., 2016; Williams *et al*., 2021). The apical portion of a graft is known as the scion and the root system is known as the rootstock (Scion:Rootstock). The capacity for two individuals to form continuous vascular connections across the graft site is known as graft compatibility (Harrison & Burgess, 1962; Moore, 1984). The inability to graft is categorized into two types of incompatibility: immediate incompatibility and delayed incompatibility (Mendel, 1936; Argles, 1937). Delayed incompatibility can present months or years after grafting, with symptoms such as swollen, over-proliferated scions, cell death in the junction, and structural instability of the stem (Eames & Cox, 1945; Moore & Walker, 1981; Andrews & Marquez, 2010). Despite a long history of grafting, humans still struggle to understand the mechanisms underlying graft incompatibility. Currently, there are only a few examples where the causes of incompatibility have been identified ( Gur, 1968; Mosse & Herrero, 1951; Moore, 1986). Although there is likely a variety of species-specific cellular mechanisms that determine compatible versus incompatible graft pairings, the presence of persistent cell death in the junction is a common symptom that is observed across diverse plant families (Eames & Cox, 1945; Moore & Walker, 1981).

Cell death can be classified into two main categories: necrosis and programmed cell death (PCD; Burbridge *et al*., 2007). Necrosis is defined as uncontrolled death and is often caused by stressors such as extreme heat, radiation, or a loss of membrane potential that is so intense that genetic processes are unable to act (Hirsch *et al*., 1997; Burbridge *et al*., 2007). In contrast, PCD is the controlled and organized process of cellular destruction (Lockshin & Zakeri, 2004).

All eukaryotes have evolved an innate immune system that is capable of detecting conserved foreign molecules during infection (Janeway et al., 2001). In plants, various elicitors such as pathogen associated molecular patterns (PAMPs) and damage associated molecular patterns (DAMPs) are perceived by membrane bound pattern recognition receptors (PRRs) facing the apoplast (Jones & Dangl, 2006; Amarante-Mendes *et al*., 2018; Steinbrenner *et al*., 2020). These molecules trigger downstream pattern triggered immunity (PTI) and signal basal defense processes such as reactive oxygen species (ROS) production (Boller & He, 2009). Alternatively, some pathogens release effector proteins, which are expressed into the symplast to modify host responses and promote infection (Stergiopoulos & de Wit, 2009). These effectors are locked in an arms race with intracellular nucleotide-binding and leucine-rich repeat receptors (NLRs), which perceive effectors or proteins modified by effectors and elicit effector triggered immunity (ETI) and PCD (Wu *et al*., 2014). Currently, no such molecules, apoplastic or symplastic, have been identified as the underlying cause of graft incompatibility, where an unknown signal from one graft partner is perceived by a protein of the other, thus leading to incompatibility.

Our previous work identified tomato and pepper as an herbaceous model for delayed incompatibility (Thomas *et al*., 2022, 2023). Despite several studies investigating pepper graft compatibility, much remains unknown about the underlying mechanism (Deloire & Hébant, 1982; Ives, 2012; Kawaguchi *et al*., 2008; Zeist *et al*., 2018). To explore this, we grafted tomato to *Capsicum annuum* varieties, Cayenne, Doux des Landes, California Wonder, and *Capsicum chinense* variety Habanero. We found that tomato-*Capsicum* heterografts are all incompatible and exhibit failed xylem reconnections, weakened stem stability, and reduced growth. Using the tomato-pepper combination with the highest graft survival rate, tomato to California Wonder, we analyzed the presence of non-viable tissue in the junction at 7, 14, and 21 days after grafting (DAG), and investigated the cause using viability staining and transcriptomics. In contrast to self-grafted controls that clear non-viable tissue from the junction, we found that incompatible grafts exhibit persistent cell death at the junction. Additionally, we utilized RNA-seq to show that incompatible grafts have a prolonged defense response following grafting, including significant upregulation of many NLRs and signaling components involved in hypersensitive response (HR). Furthermore, we identified a set of potential incompatibility marker genes that are upregulated in incompatible junctions of both tomato and pepper stems. To characterize the molecular response of incompatible grafting in relation to other biotic stress responses, we conducted a transcriptomic meta-analysis comparing the effect of grafting with pathogen infection, herbivory, and plant parasitism. We found a significant overlap in expression patterns between grafting and plant parasitism, indicating similar mechanisms underpin interspecies plant-to-plant interactions. Lastly, we identified a suite of over 1000 genes that are uniquely upregulated in incompatible grafts but not other biological stressors; among these genes, we identified genetic processes involved in immune responses and DNA damage. Together, this work supports a model in which tomato and pepper exhibit genetic incompatibility, which is potentially induced by incompatible cross-species NLRs that trigger the production of defensive compounds, upregulate programmed cell death, and eventually lead to genotoxic DNA damage. Genetic incompatibility between tomato and pepper would be the first identified instance of a hyper-immunity based incompatibility in a cross-species grafted crop.

## Materials and methods

### Plant materials and growth conditions

*Capsicum annuum* var. California Wonder (CW), RC Cayenne (Cayenne),Doux des Landes (DDL), and *Capsicum chinense* var. Habanero and *Capsicum chinense* (pepper), and *Solanum lycopersicum* (tomato) seeds were used for graft compatibility screening (Method S1). 21 day old pepper seedlings and 14 day old tomato seedlings were grafted (Method S2-3).

### Characterizing graft compatibility

30 DAG the vascular connectivity of tomato and pepper junctions was assayed using propidium iodide staining (Method S4). Graft junction integrity was tested using manual bending (Thomas *et al*., 2022). (Method S5). *Capsicum annuum* var. CW and *Solanum lycopersicum* Var. M82 were used to conduct 3-point bend tests at the University of Delaware. Structural mechanics of the graft junction were assessed by 3-point bend testing (Method S6) (Ennos *et al*., 1993; Goodman & Ennos, 2001; Hostetler *et al*., 2022).

### DAMP Assay

Hypocotyl explants from *Capsicum annuum* var. CW and *Solanum lycopersicum* Var. M82 were placed on callus-inducing media for 7 days (Method S7). The hypocotyl tissue was then placed onto media which either previously cultured tomato or pepper tissue for 7 additional days. The area of the explants was measured after 7 days on the experimental media.

### Grafting for TUNEL, trypan blue staining, and RNA-seq

*Capsicum annuum* var. (CW) and *Solanum lycopersicum* Var. M82 were grown as described above. 36 of each tomato and pepper species were left ungrafted. The rest of the plants were grafted as described above in the following combinations: 50 tomato:tomato, 50 CW:CW, 70 tomato:CW, and 70 CW:tomato. Ungrafted CW and tomato plants were included in the recovery procedure. Plastic domes were vented 7 DAG and removed 14 DAG.

### Trypan Blue staining

Stems from 7, 14, 21 DAG, and ungrafted plants were collected and stained with 1% Trypan Blue as previously reported (Method S8; Fernández-Bautista, 2016).

### TUNEL Assay

A 0.5 cm piece of the junction from 7, 14, 21 DAG, and ungrafted plants were used to image PCD. Assays were performed using the Promega DeadEnd™ Fluorometric TUNEL System (Method S9).

### Transcriptomic Analysis

*Capsicum annuum* var. CW and *Solanum lycopersicum* Var. M82 were grafted as previously described. A 0.5 cm of the junctions of 7, 14, 21 DAG, and ungrafted plants were collected from 5 biological replicates for each sample. Each piece of tissue was flash-frozen and ground with a mortar and pestle. Total RNA was purified and 3’ Seq libraries were constructed at the Cornell Institute of Biotechnology, Biotechnology Resource Center, and the libraries were sequenced on an Illumina NextSeq 500/550 using an Illumina High-output kit (Method S10). Fasta files were processed to yield raw reads and differential expression analysis was performed using DESeq2 (Method S10; Love *et al*., 2014). Putative orthogroups were determined using OrthoFinder with Diamond as the sequence search program (Method S11; Buchfink *et al*., 2014; Emms & Kelly, 2019). Publicly available RNA-seq data was downloaded and processed to yield raw read counts (Method S12).

### Statistical analysis and image analysis

All statistical computation and graph generation were performed in R v4.1.2 (R Core Team, 2021) (Method S13).

## Results

### Incompatible tomato and pepper heterografts are characterized by low survival, reduced growth, failed vascular connectivity, and physical instability

To investigate grafting between *Solanum lycopersicum* (tomato) and *Capsicum* (pepper) species, we performed a graft compatibility assay between self-and reciprocal grafts of *Solanum lycopersicum* var. M82 and *Capsicum annuum* varieties Cayenne, Doux des Landes (DDL), California Wonder (CW), and *Capsicum chinense* var. Habanero 30 DAG (Fig. 1, Fig. S1a-d, Table S1). Compared with self-grafted controls, heterografted tomato/pepper combinations exhibited significantly lower survival and higher break rates based on bend testing (Fig. 1a). Despite a low survival rate, self-grafted DDL plants that persisted formed strong graft junctions and were able to withstand the bend test (Moore, 1983; Thomas *et al*., 2022, 2023). Previous work reported DDL as a compatible graft partner with tomato, yet when we challenged the integrity of the graft using the bend test, the plants broke at the junction 92% of the time, indicating a high level of incompatibility that was previously undetected (Deloire & Hébant, 1982). We also analyzed growth of the grafted shoot and root systems to test for developmental restrictions. Compatible shoot and root systems were 92.7% and 38.2% larger than incompatible plants (Figure 1b-c, Table S2). Additionally, the scion and stock diameters 2 cm above or below the junction were significantly restricted in their lateral development in incompatible grafts compared to self-grafted controls (Kruskal-Wallis Fig. S2a-b).

**Figure 1:**
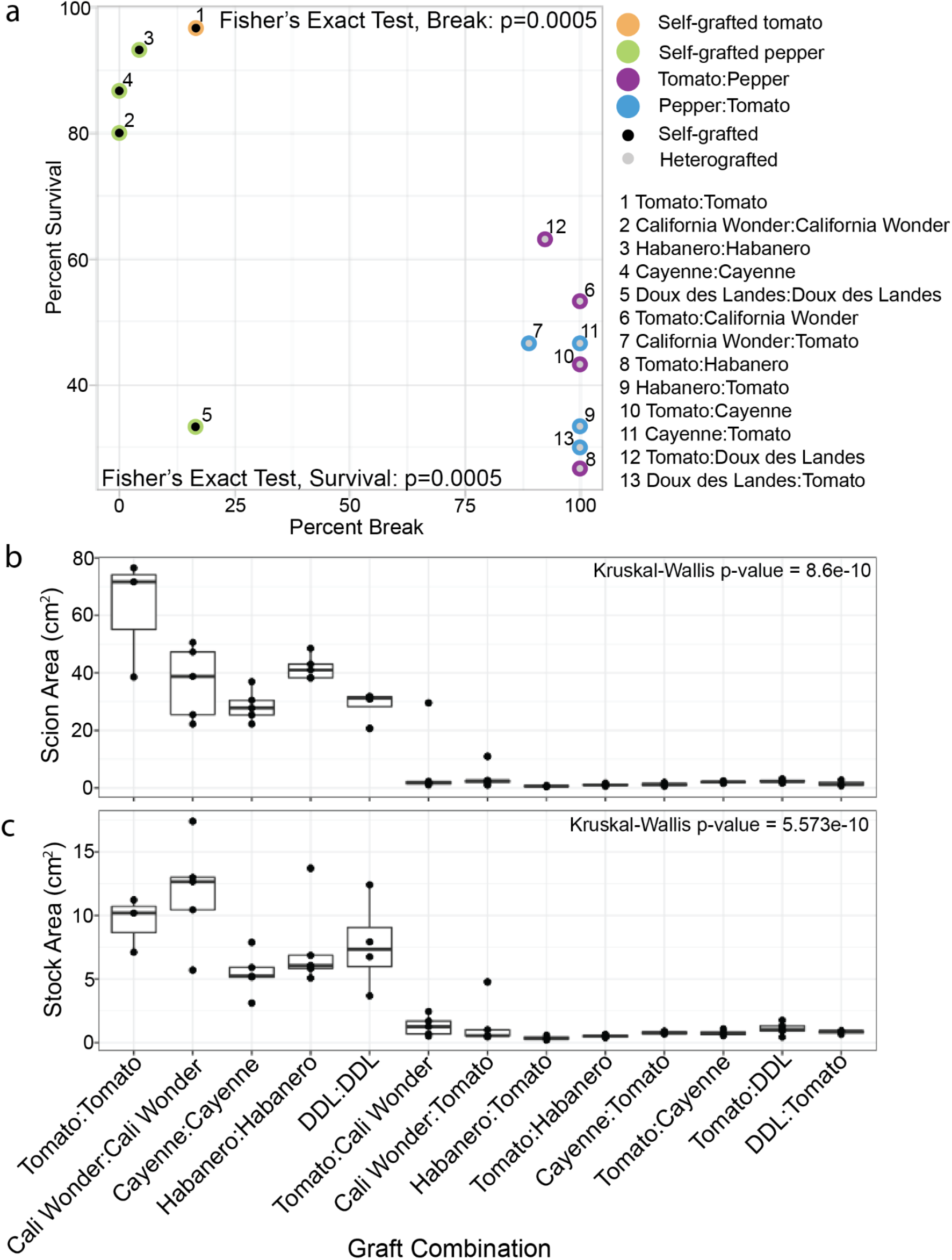
**Heterografted tomato and pepper combinations exhibit moderate survival, unstable stem integrity, and reduced growth**. The relationship between percent survival (y-axis) and percent break (x-axis) is shown for all graft combinations (a). Black dots denote self-grafts, grey dots denote heterografts. Self-grafted tomato is outlined in orange. Self-grafted pepper is outlined in green. Heterografts where the scion is tomato are outlined in purple. Heterografts where the stock is tomato are outlined in blue. The identity of each data point is labeled 1-13. Percent survival n=30; For bend test sample size see Table S1. The change in stem diameter 2cm above the graft junction between 30 and 0 DAG (scion) (b). The change in stem diameter 2 cm below the graft junction between 30 and 0 DAG (stock) (c). California Wonder abbreviated to Cali Wonder, Doux des Landes abbreviated to DDL. Biological replicates are depicted as jitter and described in Table S2. Kruskal–Wallis one-way analysis of variance was used to detect significant differences between self-and heterografted combinations. p-value <0.05.

To examine the vascular connectivity of the grafts, we analyzed the anatomical organization of junctions from every tomato/pepper combination 30 DAG (Fig. 2). Consistent with our previous findings (Thomas *et al*., 2023), all self-grafted combinations formed continuous xylem bridges across the graft junction (Fig. 2b, d, j, p, v), demonstrating compatibility (Moore, 1981; Melnyk *et al*., 2015; Thomas *et al*., 2022). Tomato grafted to any of the pepper varieties, formed non-vascular parenchymatous connections across the graft but failed to form xylem bridges (Fig. 2f, h, l, n, r, t, x, z). We noticed that overproliferated callus (Fig. 2l,t,x), as well as adventitious root growth (Fig. 2f, n, t) were common features in these incompatible combinations.

**Figure 2:**
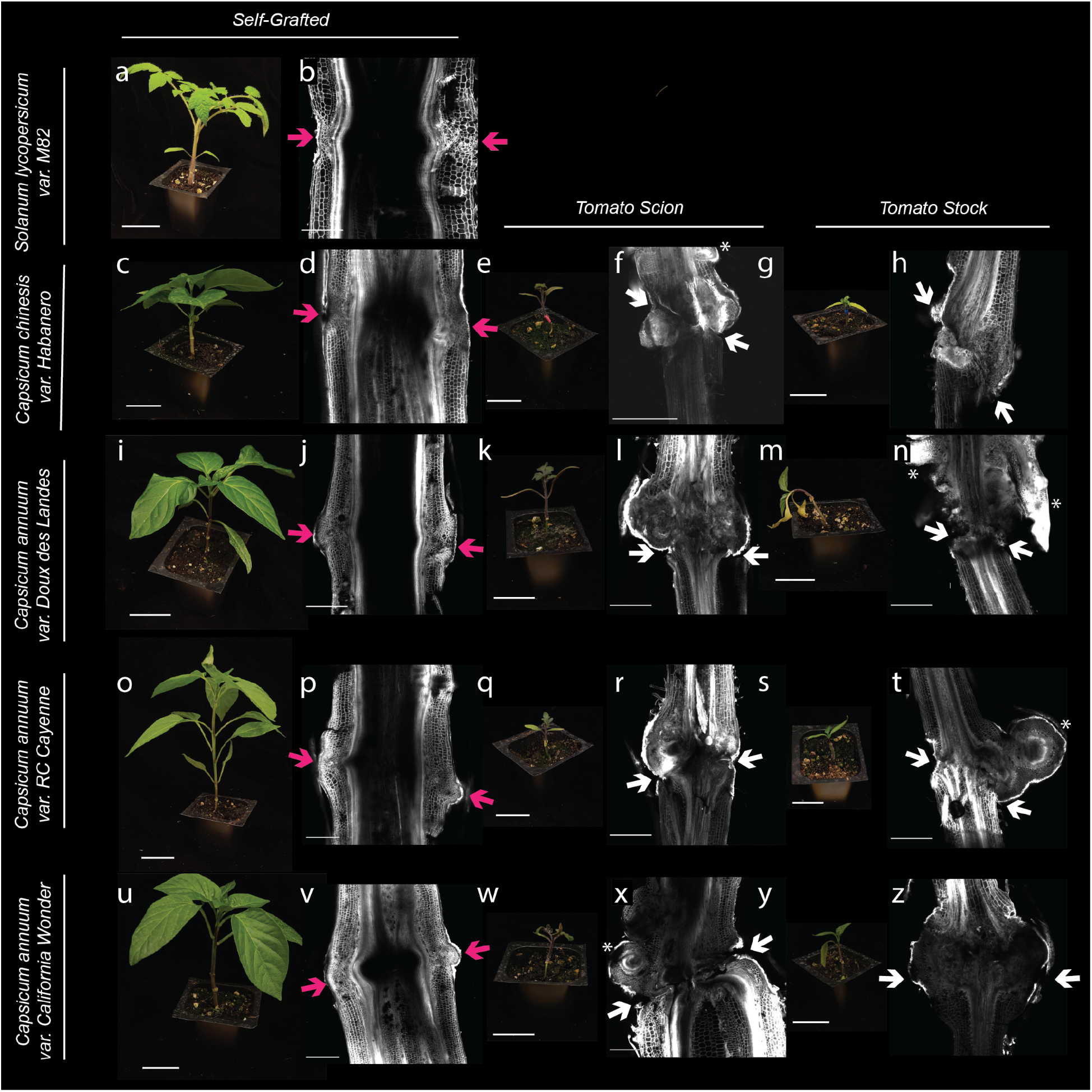
Heterografted pepper fails to form vascular connections and shows a significant decrease in size 30 DAG. (a, c, e, g, i, k, m, o, q, s, w, u, w, y) Representative photographs and (b, d, f, h, j, l, n, p, r, t, v, x, z) confocal micrographs for self-grafted tomato (a-b), self-grafted habanero (c-d), tomato:Habanero (e-f), Habanero:tomato (g-h), self-grafted Doux des Landes (DDL) (i-j), tomato:DDL (k-l), DDL:tomato (m-n). self-grafted Cayenne (o-p), tomato:Cayenne (q-r), Cayenne:tomato (s-t), self-grafted California Wonder (CW) (u-v), tomato:CW (w-x), CW:tomato (y-z). Graft junctions were stained with propidium iodide and imaged on a confocal microscope. Pink arrows indicate a successful graft junction with a healed xylem, white arrows indicate a failed vascular reconnection and white Asterix highlight adventitious roots. All plant images have scale bars are 5 cm, and all micrograph scale bars are 1000 µm.

Because CW exhibited high survival rates when heterografted with tomato, we selected this genotype for further analysis. Incompatible grafts are commonly discovered when the junction breaks, due to failed vascular connectivity or cell death in the junction (Eames & Cox, 1945; Moore & Walker, 1981; Andrews & Marquez, 2010). With a better understanding of the vascular anatomy of compatible and incompatible plants, we sought to determine if we could quantify the instability observed in the manual bend test using a quantitative 3-point bend test. Congruent with reduced biophysical stability observed with the 3-point bend test, we found that there was a significant reduction in the structural stiffness of the heterografted junctions compared to self-and ungrafted stems (Fig. S1e-f, Table S3).

### Incompatible graft junctions accumulate significantly more non-viable tissue than compatible grafts

Cell death is a common symptom associated with incompatible grafts (Moore, 1983). To examine the extent to which tomato/pepper (CW variety) heterografts exhibit elevated levels of cell death, we collected tissue from ungrafted tomato and pepper, self-graft tomato and pepper, and reciprocally heterografted tomato and pepper at 7, 14, and 21 DAG (Fig. S3). To quantify cell death in the junction, we used trypan blue staining to detect regions of deep tissue cell death (Fig. 3, Table S4). We measured a 2.5 mm region of the junction, including any callus present at the interface. When we considered just this sample area, the area of all graft junctions increased at a similar rate, independent of the stem diameter (Fig S5a). To quantify the percent of non-viable tissue (NVT) versus viable tissue, we made a macro in ImageJ to extract tissue that was deeply stained with trypan blue (Fig. 3ak, Fig. S4) and divided this area by the entire area of the junction (Fig S5b). We first analyzed ungrafted tomato (Fig. 3a, g, m) and pepper (Fig. 3b, h, n) stems that were the same age as the grafts we harvested at 7, 14, and 21 DAG. At most, the ungrafted stems from tomato and pepper contained 0.341% and 0.147% NVT respectively. Self-grafted tomato graft junctions consisted of 13.0% NVT at 7 DAG, which decreased to 3.51% by 21 DAG (Fig. 3u, aa, ag; Wilcoxon Paired Test p = 0.0589). Similarly, self-grafted pepper junctions contained 24.8% NVT at 7 DAG but steadily decreased to only 2.92% by 21 DAG ( Fig. 3v, ab, ah; p = 2.78E-02). Unlike the self-grafts, which exhibited decreasing NVT over time, tomato:pepper and pepper:tomato incompatible grafts maintained a consistent percent of NVT over the three week sample period (Fig. 3s-aj). Tomato:pepper junctions contained 20.9%, 20.2%, and 20.8% NVT at 7, 14, and 21 DAG, respectively (Fig. 3w, ac, ai), and reciprocal pepper:tomato junctions exhibited similar levels of NVT: 21.9%, 17.9%, and 17.1% NVT at 7, 14, and 21 DAG, respectively (Fig. 3x, ad, aj). Overall, tomato and pepper exhibited prolonged cell death up to three weeks post-grafting relative to self-grafted controls.

**Figure 3:**
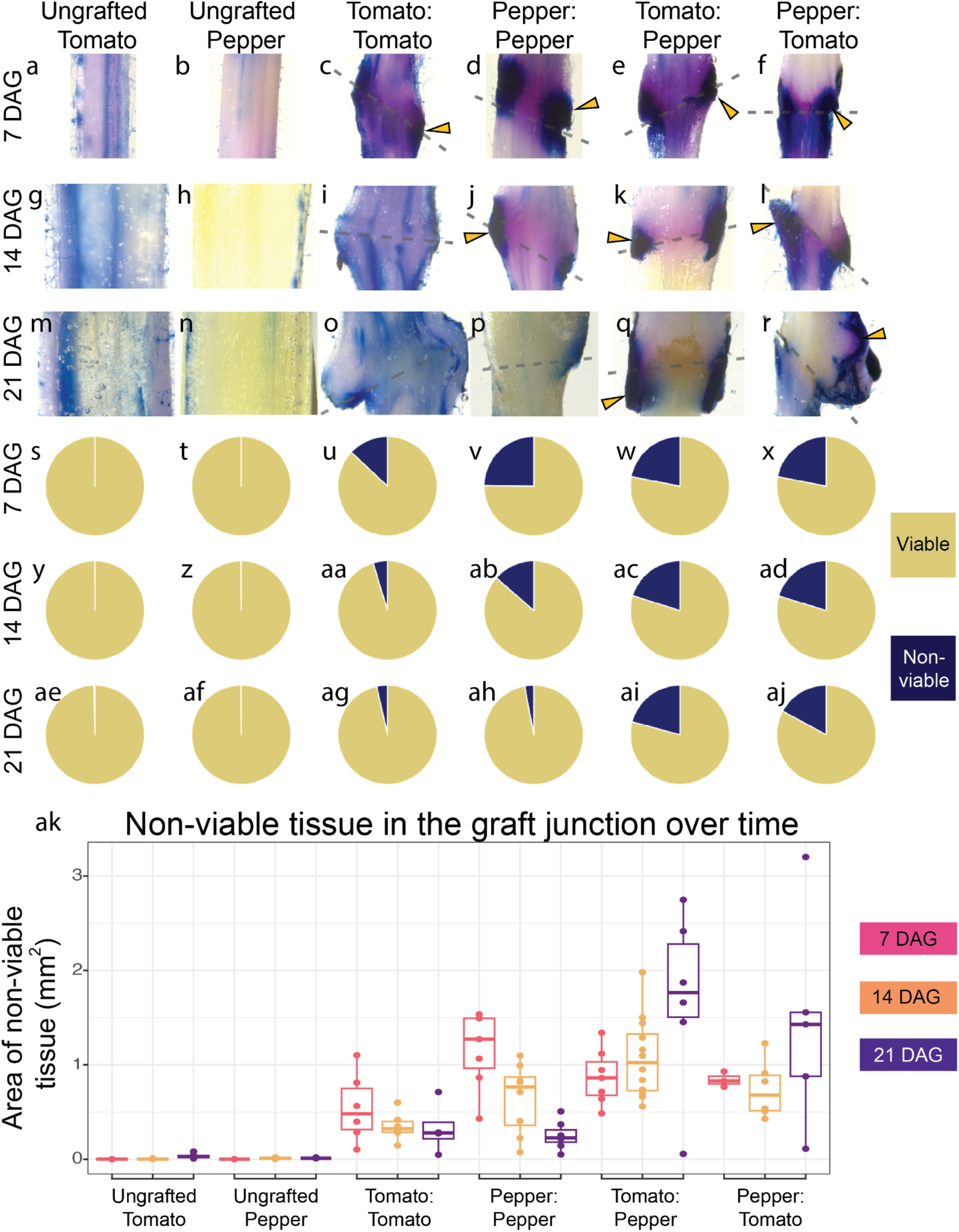
Incompatible grafts contain persistent nonviable tissue over time. (a-r) Representative images of 2.5 mm long graft junctions at 7, 14, and 21 DAG stained with Trypan Blue. A representative ungrafted tomato stem and the percent of non-viable tissue (NVT) are shown at 7 DAG (a, s), 14 DAG (g, y), and 21 DAG (m, ae). A representative ungrafted pepper stem and the percent of NVT at 7 DAG (b, t), 14 DAG (h, z), and 21 DAG (n, af). A representative self-graft tomato junction and the percent of NVT at 7 DAG (c, u), 14 DAG (i, aa), and 21 DAG (o, ag). A representative self-grafted pepper junction and the percent of NVT at 7 DAG (d, v), 14 DAG (j, ab), and 21 DAG (p, ah). A representative tomato:pepper junction and the percent of NVT at 7 DAG (e, w), 14 DAG (k, ac), and 21 DAG (q, aI). A representative pepper:tomato junction and the percent of NVT at 7 DAG (f, x), 14 DAG (l, ad), and 21 DAG (r, aj). Yellow arrows point to examples of deep tissue death; dashed lines signify the graft site; all junctions are 2.5 mm tall (a-r). (s-aj) The percent of cell death and (ak) the area of cell death in the junction of all graft combinations at 7, 14, and 21 DAG. Pink boxplots are 7 DAG, orange boxplots are 14 DAG, and purple boxplots are 21 DAG. Biological replicates are depicted as jitter (ak) as well as described in detail in Table S4.

Previous work has attributed this incompatible symptom of NVT to the accumulation of trapped cellular debris that creates a necrotic layer in the graft (Tiedemann, 1989). However, the active accumulation of NVT through programmed cell death (PCD) provides an alternative explanation. To test whether NVT accumulation in incompatible grafts was due to PCD, we performed terminal deoxynucleotidyl transferase dUTP nick end labeling (TUNEL) assays on graft junctions from all graft combinations at 7, 14, and 21 DAG (Fig 4, Fig S6-S7). We observed that ungrafted tomato and pepper stems contain cells undergoing PCD at low rates within the stem tissues particularly in xylem and epidermal cells, and indeed PCD could be detected in graft junctions as well (Fig S6-7). Despite detecting PCD in the peripheral areas of the graft junction that overlapped with the region of NVT identified by trypan blue, the high amount of developmental cell death due to vasculogenesis confounded our ability to quantify differences in PCD between compatible (Fig. 4a,b,e,f) and incompatible (Fig. 4c,d,g,h) grafts.

**Figure 4:**
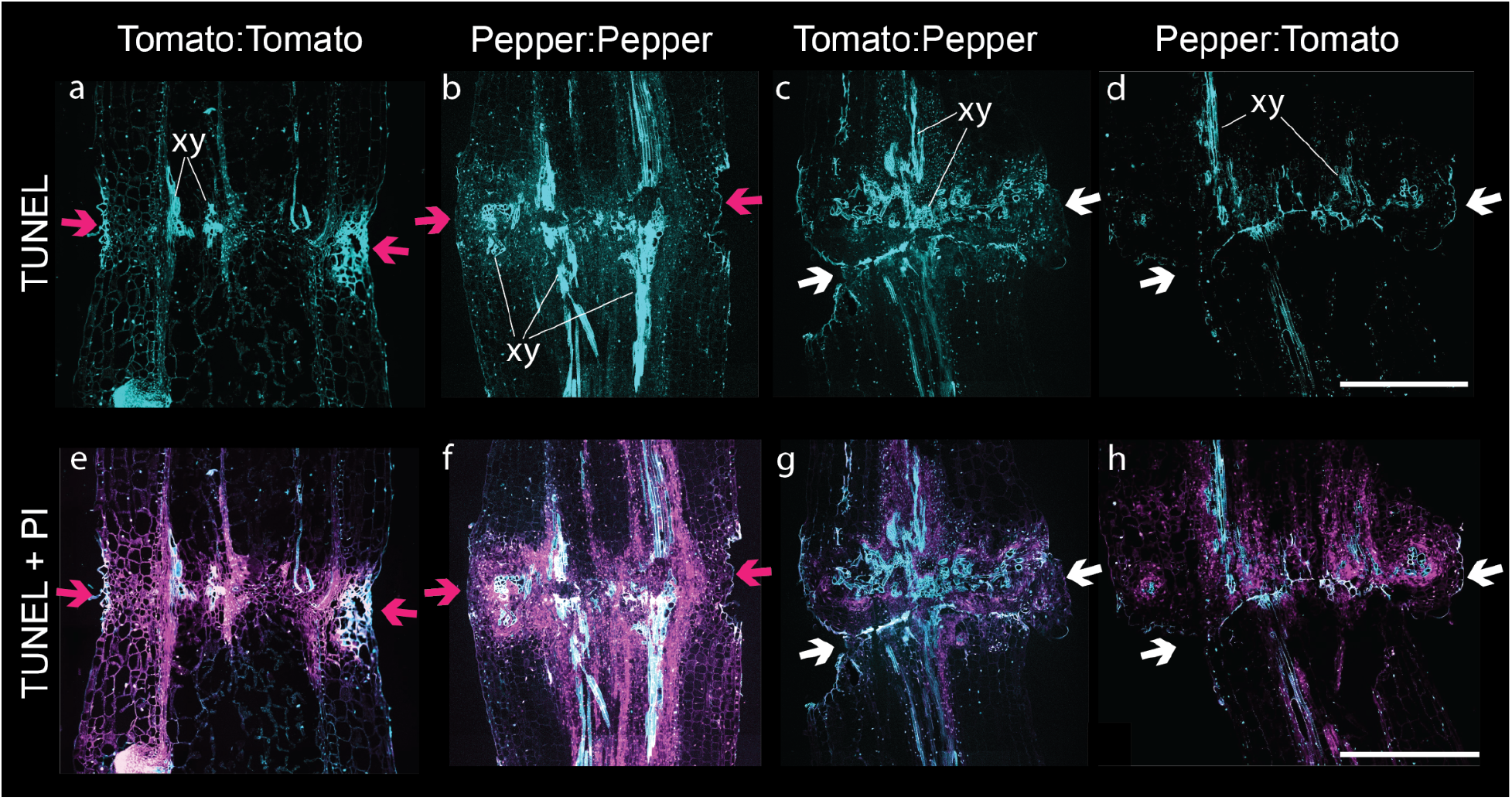
Developmental programmed cell death is present in all graft junctions regardless of compatibility. A representative graft junction from (a,e) tomato:tomato, (b,f) pepper:pepper, (c,g) tomato:pepper, (d,h) pepper:tomato 14 DAG. (a-d) TUNEL fluorescein-12-dUTP-labeled DNA and autofluorescence are false-colored cyan. (e-h) the TUNEL fluorescence merged with propidium iodide (false-colored magenta) staining nucleic acid and cell walls. Pink arrows indicate a successful graft junction with healed xylem, and white arrows indicate a failed vascular reconnection. Examples of newly developed xylem are labeled (xy). All images are equal and the scale bar is 500 µm.

To understand the cause of the NVT present in the incompatible grafts, we first investigated if DAMPs, which activate PAMP-triggered immunity (PTI) upon cellular rupture during infection and herbivory, also play a role in determining incompatible species combinations (Brutus *et al*., 2010; Ferrari *et al*., 2013; Nothnagel *et al*., 1983). To see if a component of the cell wall could act as an antagonist to inter-species grafting, we designed an *in vitro* assay to test for DAMP-induced changes in callus growth similar to work conducted in quince (Moore, 1986). Tomato and pepper explants were allowed to grow on media containing either tomato or pepper wound exudates for 7 days. If wounding caused the secretion of an inhibitory chemical during callus formation, the hypocotyls growing on cross-species exudates would be stunted. Despite a significant difference in overall growth rates between tomato and pepper explants, the presence of cross-species DAMPs had no effect (Fig. S8, Table S5). Our results indicate that cross-species secreted exudates do not affect callus growth; however, this does not rule out the role of DAMPs in triggering graft-incompatibility during other stages of junction formation.

### Tomato and Pepper heterografts express prolonged transcriptional defense profiles

To further investigate the underlying cause of NVT in incompatible grafts, we collected and performed RNA-sequencing on ungrafted, self-grafted, and heterografted tissue at 7, 14, and 21 DAG (Table S6-7). When compared to ungrafted stems, self-grafted junctions expressed 4.5x, 5x, and 15x less differentially expressed genes compared to heterografts at 7, 14, and 21 DAG, respectively. (Fig. 5, Table S8-9). The reduced number of differentially expressed genes in self-grafts correlates with the healing timeline, where compatible tomato and pepper self-grafts heal within the first week (Thomas *et al*., 2022).

**Figure 5:**
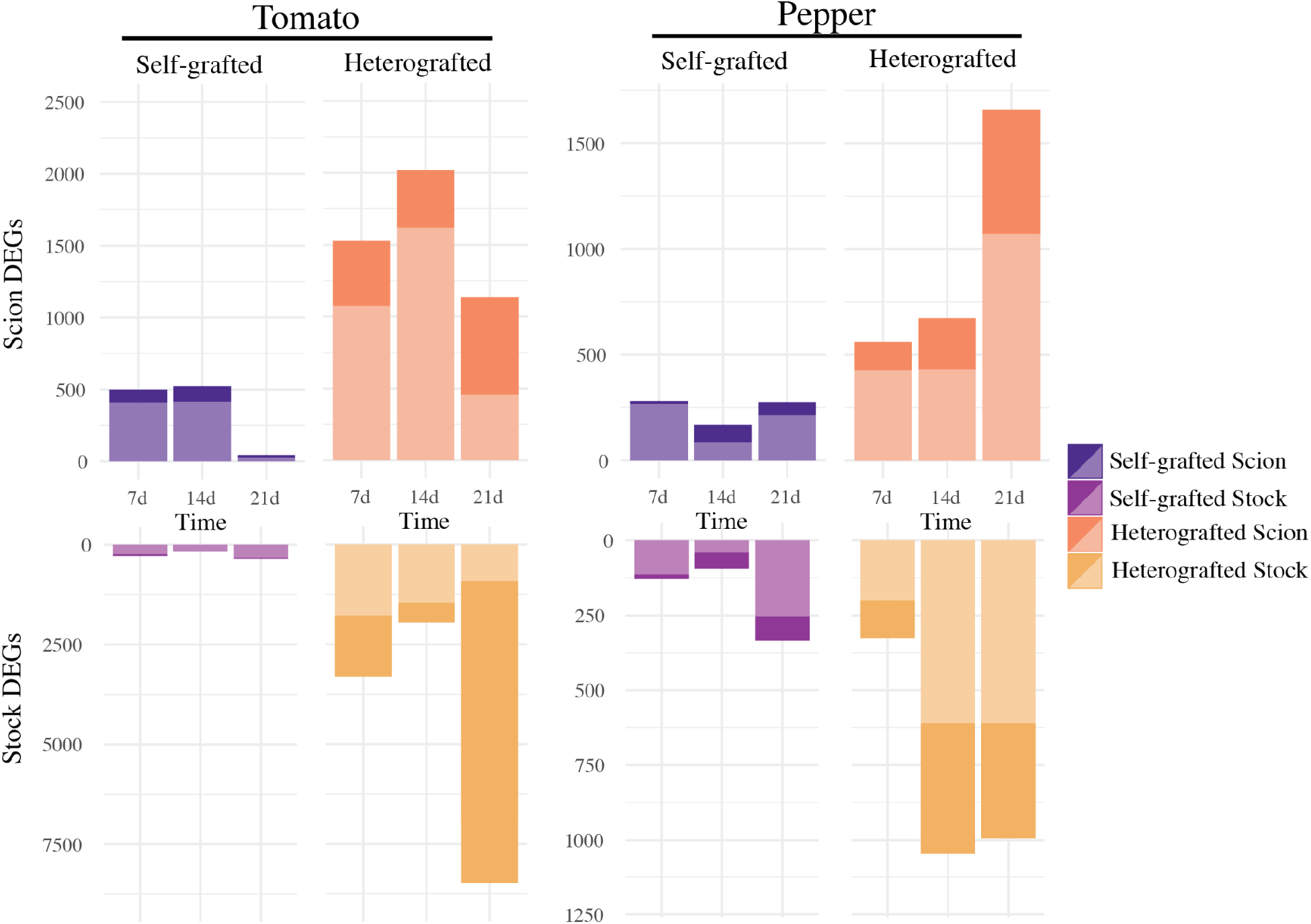
Incompatible heterografts have prolonged differential gene regulation compared to self-grafts. Differentially expressed genes (>1.5 or <-1.5, p-value<0.05) of each grafted tissue (compared to ungrafted) at each time point for tomato and pepper. Upregulated genes are shown in light colors and downregulated genes are shown in dark colors. Self-grafted scions are dark purple, self-grated stocks are light purple, heterografted scions are orange, and heterograft stocks are yellow. Each combination has 3-5 bio-replicates.

To identify genes uniquely upregulated in the incompatible grafts, we used likelihood ratio testing (Table S10). Distinct genes were expressed in the scion and stock, with only a fraction in common at any time point, suggesting that the genetic response in incompatible grafts is spatially and temporally regulated (Fig. S9a-f). For example, at 7 DAG, 1530 and 2380 genes were uniquely upregulated in the scion and stock of incompatible tomato grafts (Fig. S9). Of these 3910 genes, only 576 were shared between scion and stock. The percent of genes upregulated in the scion that were also upregulated in stock for tomato was only 38%, 2%, and 11% of the total scion DEGs at 7, 14, and 21 DAG respectively. Similarly, genes upregulated in the pepper scion that were also upregulated in the stock made up only 6%, 29%, and 17% of the total scion genes at 7, 14, and 21 DAG. Additionally, scion tissue shared more genes across time than stocks, further supporting that the position in relation to the graft junction, holds a significant role in the genetic process (Fig. S9g-j).

Using significantly upregulated genes from either tomato:pepper or pepper:tomato incompatible graft combinations, we performed GO term enrichment (Figure S10, Table S11). We found that processes associated with defense and stress displayed the highest enrichment in heterografted tomato stocks 14 DAG and pepper stocks 7 DAG (Fig. 6a-b). To further explore how defense processes might be involved in the incompatible response, we targeted NLRs and downstream molecular signaling involved in defense for deeper analysis. Using a collection of 320 previously annotated tomato NLRs (Bashir *et al*., 2022), we identified 97 defense-related receptors that were significantly upregulated in incompatible grafts (Fig. 6c, Table S12). Of these 97 NLRs, 82 were upregulated in the pepper:tomato 14 DAG sample, indicating this is a critical time point for activating defense-related molecular responses to incompatibility. Similarly, 145 of the 356 annotated pepper NLRs were upregulated during incompatible grafting (Fig. 6d), with a pronounced molecular signature of 101 NLRs upregulated in tomato:pepper grafts 7 DAG (Lee *et al*., 2021). Notably, stock tissue from both tomato and pepper incompatible grafts exhibit highly upregulated NLR expression within the first two weeks post-grafting (Fig 6a-b). In the absence of an effector protein or pathogen, overexpression of NLRs can trigger autoimmunity that leads to hypersensitive response (HR; Freh *et al*., 2022). To test whether cell death in incompatible grafts could result from HR, we analyzed the expression of tomato and pepper orthologs EDS1, SAG101, and PAD4, which are known regulators of HR in arabidopsis (Fig. 6e, Table S12; Rietz *et al*., 2011; Zhu *et al*., 2011; Gantner *et al*., 2019). Again, we identified incompatible graft-specific upregulation, especially at 14 DAG, for these HR regulators (Fig. 6e).

**Figure 6.**
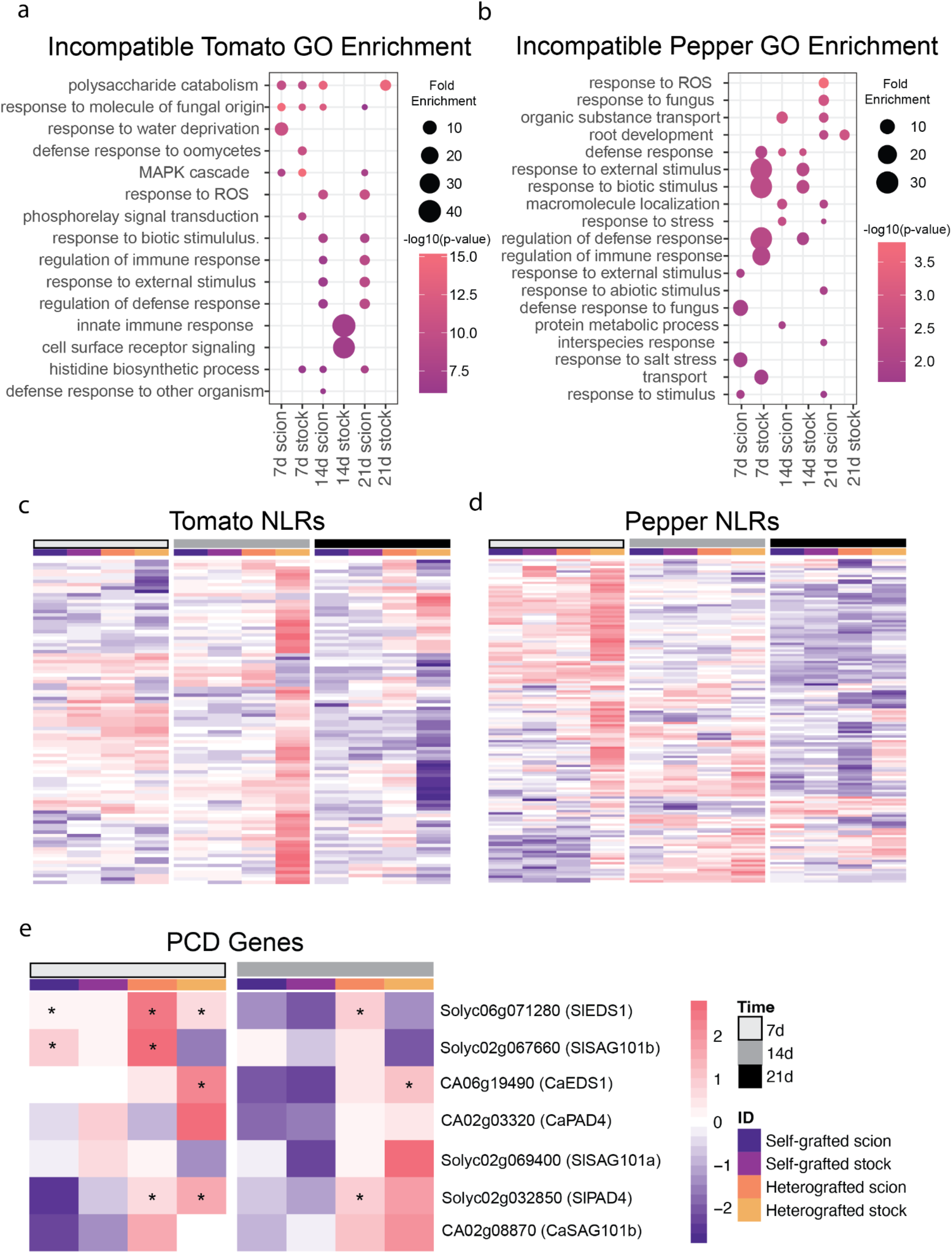
**Heterograft-specific upregulated genes are involved in defense response. (**a-b) Uniquely upregulated heterografted genes were determined by performing likelihood ratio testing (p<0.05) on ungrafted, self-graft scion, and heterografted scion as well as ungrafted, self-grafted stock, and heterografted stock tissue. The genes upregulated in only the heterograft tissue were used to perform GO enrichment. GO terms enriched in heterografted tomato tissue at 7, 14, and 21 DAG (a). GO terms enriched in heterografted pepper tissue at 7, 14, and 21 DAG (b). (c-d) Log-fold change of NLRs in grafted tissue compared to ungrafted tissue of tomato (c) and pepper (d). (e) The log-fold change of genes involved in hypersensitive response in grafted vs. ungrafted tissue. The log-fold change was scaled by row. The tissue is denoted by the colored columns where self-grafted scions are dark purple, self-grafted stocks are light purple, heterografted scions are orange, and heterografted stocks are yellow. The days after grafting were denoted by colored columns where 7 DAG are white, 14 DAG are grey, and 21 DAG are black. Asterisks denotes p-value<0.05 and log-fold change greater than |1.5|.

Next, to explore the role of hormonal regulation in graft compatibility, we identified the closest tomato and pepper putative homologs for annotated Arabidopsis genes involved in salicylic acid (SA), jasmonic acid (JA), and ethylene biosynthesis and response (Fig. S11a-c, Table S12) All three of these hormones are known to play a role in defense processes, with SA serving a critical function in NLR-induced HR (Enyedi *et al*., 1992; Lorenzo *et al*., 2003; Koornneef & Pieterse, 2008). In addition, JA and ethylene are associated with graft junction formation (Wang *et al*., 2020; Thomas *et al*., 2022). At 7 DAG, SA, JA, and ethylene biosynthesis were upregulated in self and incompatible grafted samples relative to ungrafted controls, indicating that the 7 day response is dominated by general graft healing processes. By 14 and 21 DAG, SA, JA, and ethylene biosynthesis and perception were predominantly upregulated in the incompatible scions compared to self-grafted controls (Fig S11). Congruent with this response, we noticed that the tomato and pepper orthologs for PR1, a defense gene downstream of SA signaling, was upregulated at 21 DAG in incompatible scions. The expression of these genes three-weeks after grafting indicates that the prolonged incompatible graft response is related to defense processes which may be activated or mediated by SA, JA, and ethylene hormonal pathways.

Another hormonal-regulated defense process that was significantly enriched across our incompatible graft time points is the biosynthesis of steroidal glycoalkaloids (SGAs; Fig. S12, Table S12). SGAs are a class of jasmonate-dependent defensive compounds produced by Solanaceous species (Cárdenas et al. 2016; Milner et al. 2011; Itkin et al. 2013; Panda et al. 2022). Upon further investigation, we were able to find that many genes in SGA biosynthesis (GAME1,4,6,7,11,12,17,18, and MKB1) were significantly upregulated in the incompatible tissue, especially in the scion (Nakayasu *et al*., 2018). Since this response is shared in tomato and pepper, it is possible that SGA biosynthesis could be triggered by the graft incompatibility immune response (Abdelkareem et al. 2017). Furthermore, SGA content could be a useful metric for gauging graft compatibility in Solanaceae.

We also explored molecular markers for ROS production, which is capable of causing cell death. We examined the expression profiles of known RBOHs (Li *et al*., 2019; Raziq *et al*., 2022), and found that of the 8 RBOHs annotated in tomato, only SlRBOH1 and SlRBOHF were both upregulated in incompatible tissue (Fig. S11d, Table S12). Surprisingly, none of the antioxidant enzymes previously shown to be upregulated alongside RBOHs were significantly upregulated in any of our incompatible grafts (Raziq *et al*., 2022). This data indicates that ROS production is not a dominant by-product of tomato-pepper graft incompatibility at 7 DAG and beyond.

We observed that the stocks of incompatible grafts exhibited high levels of RNA degradation, especially at 21 DAG (Fig. S13). RNA degradation over time in incompatible tissue might be a by-product of genotoxic stress or DNA damage as a part of NLR-activated HR (Rodriguez *et al*., 2018; Nisa *et al*., 2019). To see if known genes associated with PCD in Arabidopsis could shed light on the genetic response seen in the incompatible tissue, we identified putative orthologs for programmed cell death indicator genes from Arabidopsis (Olvera-Carrillo *et al*., 2015). We found that several orthologs for PCD-associated genes were upregulated in incompatible grafts, including HSFB1, ATHB12, and LEA7. Although our TUNEL assays were inconclusive due to the confounding effects of vasculogenesis during graft formation, this data provides molecular support for the role of PCD in promoting persistent cell death in incompatible tomato-pepper graft junctions.

Another interesting group of genes uniquely upregulated in the incompatible grafts were identified plant paralogs to BREAST CANCER SUSCEPTIBILITY GENE 1 (BRCA1, Solyc09g066080, Solyc12g041980) and BRCA1 ASSOCIATED RING DOMAIN PROTEIN 1 (BARD1, Solyc05g016230). In mammals, these genes form a homodimer which is required for homologous recombination, a mode of DNA repair following genotoxic stress. Homologous recombination is a less common method of DNA-repair in eukaryotes, whereas non-homologous end joining is the most prevalent mode. Regardless, the Arabidopsis paralogs for BRCA1/BARD1 are upregulated by genotoxic damage, so it is possible that the NLR-induced immune response in incompatible grafts leads to DNA damage which triggers BRCA1/BARD1. This is especially interesting considering that incompatible tissue showed a genetic overlap with NLR-induced HR, increased defensive processes, an indication of reduced DNA quality over time, and prolonged NVT in the junction. Together this suggests that incompatible grafts might indeed undergo a type of cross-species immune response inducing DNA breakdown and subsequently leading to cell death at the graft junction.

### Evolutionary conservation of incompatible tomato and pepper gene families

To determine whether incompatible graft response genes were conserved between tomato and pepper genomes, we generated strict orthogroups between tomato, pepper, and Arabidopsis (Table S14), and then identified significantly upregulated genes (between grafted versus ungrafted controls) with shared ortholog groupings in both tomato and pepper (Tabel S15-16). For instance, out of the 1074 and 428 genes upregulated in the heterografted scions of tomato:pepper and pepper:tomato at 7 DAG, there were 69 orthogroups conserved between the two species. We identified relatively more shared orthogroups in incompatible graft samples versus self-grafted controls, especially with respect to incompatible scion samples (Fig. 7a). Even when considering the magnitude of DEGs between samples, the number of shared orthogroups in incompatible tissue remains proportionately higher than self-grafted tissue. This finding indicates that molecular responses to incompatibility share a high degree of overlap between the tomato and pepper genomes. From this analysis, we identified ERF114 (Solyc03g118190/CA03g31320) as a shared orthogroup that is present in incompatible grafts at 7, 14, and 21 DAG. ERF14 is closely related to RAP2.6L (RELATED to AP2.6L; Fig. 7b), a wound-responsive transcription factor that exhibits overlapping expression with auxin depletion and high levels of JA in stock tissue within the first 24 hours of grafting (Asahina *et al*., 2011; Matsuoka *et al*., 2018; Lakehal *et al*., 2020). Although AtERF114 has not been tested for a direct role in grafting, similar to RAP2.6L, it has been shown to be upregulated under high JA (Lakehal *et al*., 2020). Incompatible-specific expression of SlERF114 could be explained by its role in ectopic xylem and lateral root formation in arabidopsis (Canher et al. 2022). This hypothesis is supported by the formation of unorganized overproliferated xylem tissue in the incompatible grafts, many of which produce adventitious roots (Fig. 2). These orthologs, much like SGA biosynthesis, serve as candidate markers for detecting incompatibility in Solanaceae.

**Figure 7.**
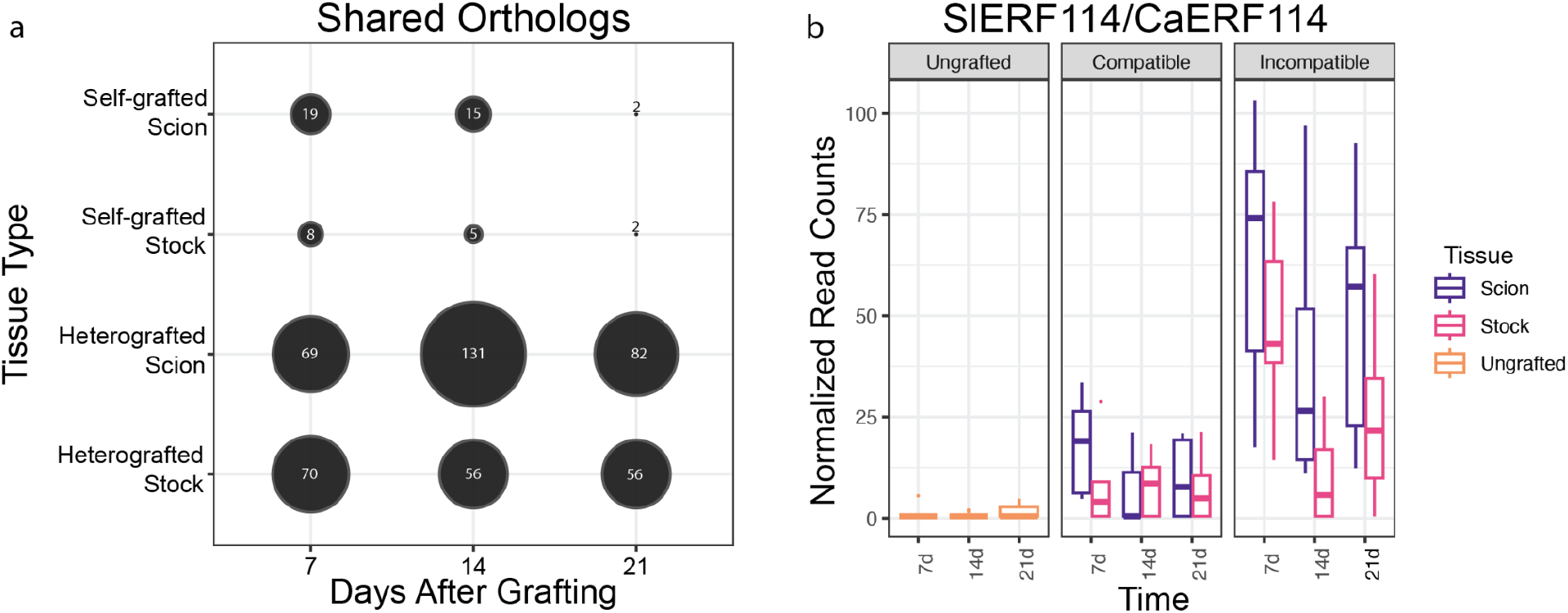
Heterografted plants share many differentially expressed putative orthologs such as ERF114. (a) Putative orthologs upregulated at any given tissue/time point in both tomato and pepper. Orthogroups were determined between *Solanum lycopersicum, Capsicum annum,* and *Arabidopsis thaliana* using OrthoFinder, where each gene corresponded to an orthogroup. Upregulated genes for all graft combinations were determined in comparison to ungrafted stems. A shared ortholog was determined if upregulated genes (lfc >1.5, p-value<0.05) from both tomato and pepper at a common tissue/time point were linked to the same orthogroup. (b) Normalized read counts of SlERF114 and CaERF114 across time. Read counts for tomato and pepper were normalized, combined, and faceted by tissue type. Boxplot color denotes tissue origin; Ungrafted tissue is orange, scion tissue is purple, and stock tissue is pink.

### Incompatible grafting upregulates a set of unique defense processes

Our analyses of incompatible graft responses indicate that both tomato and pepper upregulate strong disease resistance-related molecular responses. To test whether this response is specific to incompatible grafting, or whether these genes share overlapping functions with plant immunity and defense, we compared upregulated incompatible grafting genes with published datasets of three biotic stressors: early plant-parasitism (Jhu *et al*., 2021), insect herbivory (Ke *et al*., 2021), and established necrotrophic fungal infections (47 hours post-inoculation; Srivastava *et al*., 2020). We also used an arbuscular mycorrhizal symbiosis dataset as a control for non-destructive biotic processes (Zeng *et al*., 2023). Despite these processes occurring in differing tissues and developmental stages, all datasets were moderately correlated with all 7 DAG tomato samples (Spearman Rank average correlation; 0.58; Fig. 8a) and we were able to identify shared transcriptional responses with grafted plants (Fig. 8d-g), as well as between different biotic treatments (Fig. 8b, Table S17). Additionally, all stressors were found to have a significant representation factor (RF) greater than 1 with self and heterografted tissue; meaning that there was a significantly increased overlap of genes upregulated in the stressed tissue and the grafted tissue than expected by chance (Fisher’s Exact Test with hypergeometric probability; Table S18). This is in comparison to the Arbuscular mycorrhizal fungi (AMF) dataset, which contained 1385 significantly upregulated genes but lacked enriched overlap with self-or heterografted tissue (Fig.8d)

**Figure 8:**
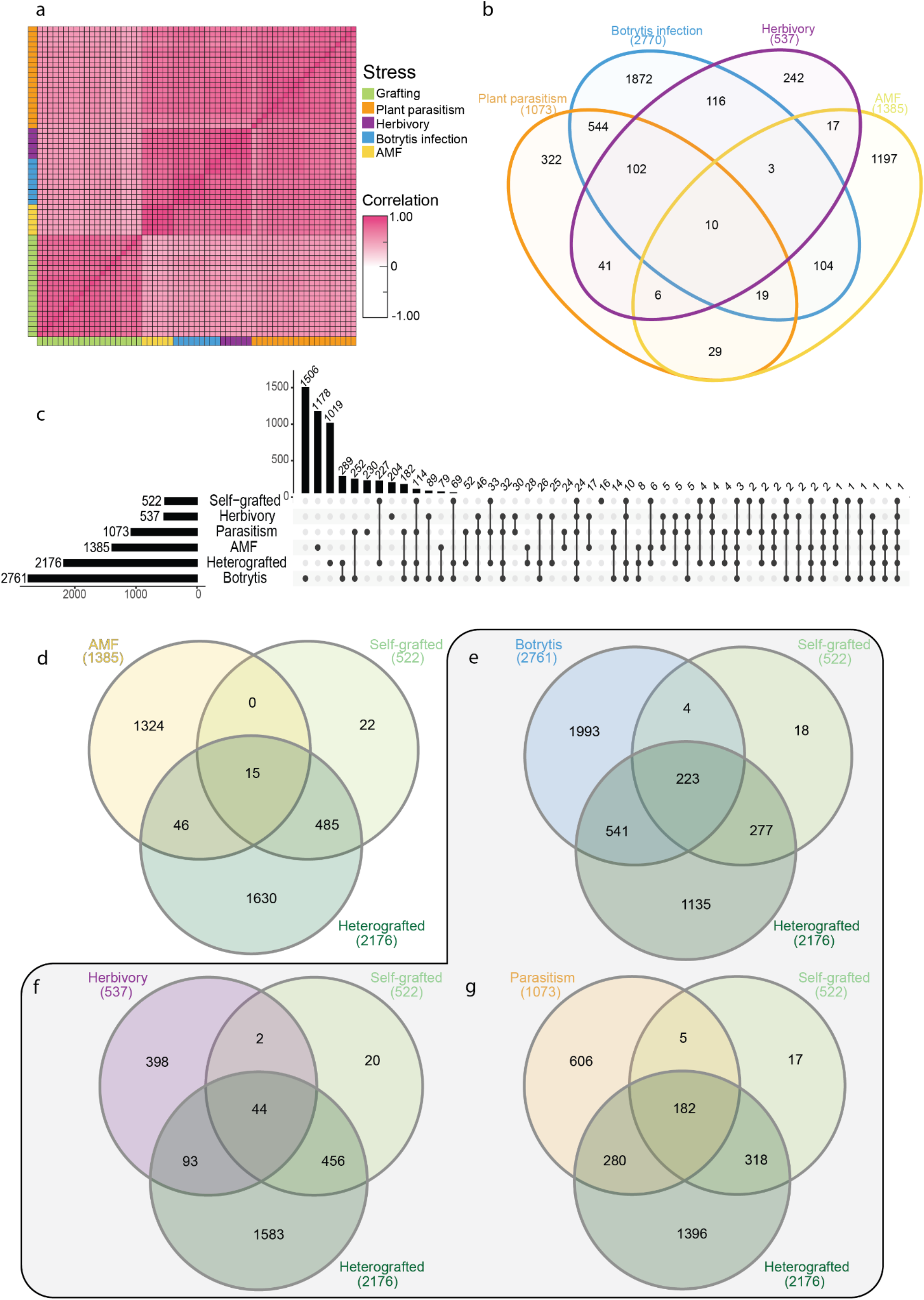
Grafting elicits unique and shared genetic processes with other biological stressors. (a) Spearman Rank Correlation between 7 DAG samples, botrytis infection, herbivory, plant parasitism, and arbuscular mycorrhizal fungi (AMF) colonization. (b) Overlap of upregulated genes from four biological processes investigated. (c) Upset plot showing the overlap between upregulated genes from the biological processes: AMF, plant parasitism, insect herbivory, and fungal infection, scion or stock self-, and scion or stock heterografted tissue 7 DAG. (d-g) Overlap of upregulated genes from scion or stock of self-or hetero-grafted tissue at 7 DAG and all biological processes. (d) The overlap between grafting and AMF, (e) botrytis fungal infection, (f) herbivory, (g) and plant parasitism. The grey outline denotes the biological stressors, whereas AMF was used as a control.

Amongst the three biotic stressors analyzed, the necrotrophic fungi, *Botrytis cinerea*, elicited the highest transcriptional responses with 2761 differentially upregulated genes. 223 of these genes were also upregulated in self and incompatible grafts. 541 genes were upregulated in only infected and incompatible grafted tissue (RF: 2.2, Fig. 8e). Shared genes were involved in defense-related processes such as RLKs, MAP kinases, LRR-proteins, and cell death, such as HSR4 (Solyc02g062550; Table S19; <u>Zhang et al.</u> <u>2014)</u>. Plants stressed with herbivory by the tobacco hornworm (*Manduca sexta*) expressed 537 upregulated genes (Fig. 8f). Of these, 44 were shared between herbivory, self-and incompatible grafted plants, 2 were uniquely shared with self-grafted datasets (RF:5.9), and 93 were uniquely shared with incompatible grafts (RF:1.9).

The parasitic plant, *Cuscuta campestris,* led to 1073 upregulated DEGs, of which 182 are shared between parasitized, self-, and incompatible grafted tissue (Fig. 8g). 280 genes were both upregulated in only parasitized tissue and incompatible grafts (RF:3.2), while 5 were upregulated in both parasitized and self-grafted, but not incompatible grafts (RF:6.4). The developmental and anatomical processes of parasitic haustorium formation and graft formation share strong parallels; both structures involve tissue reunion and the patterning of newly formed vascular connections. Given these parallels, we hypothesized that parasitism would have the greatest overlap in DEGs with grafted stems, which we found to be especially true for compatible grafts (total gene overlap, RF:5.7). Surprisingly, parasitism also shared the most significant overlap with incompatible grafts out of all 3 biotic stress treatments (total gene overlap, RF: 3.4). Enriched processes in parasitized, self-and heterografted plants include: polysaccharide catabolic processes, response to molecules of fungal origin, defense response to other organisms, and cellular response to oxygen-containing compounds. Genes from these categories include Pathogenesis-related (PR) genes, endochitinases, chitinases, and ethylene biosynthesis components. Genes upregulated in both parasitized tissue and incompatible grafts were enriched for GO terms such as MAPK cascades, regulation of defense responses, regulation of immune response, and defense response to other organisms suggesting that both incompatible grafting and plant parasitism elicit interspecies defense responses.

While we identified a significant overlap between self-, incompatible grafts, and tissue subjected to the three biotic stressors, we also identified a large set of genes that were uniquely upregulated in grafted samples only (Fig.8c). Self-and incompatible grafts uniquely upregulated 227 genes, and incompatible grafts alone expressed a unique signature of 1019 upregulated genes. These genes were enriched for GO terms including polysaccharide catabolic process and anthocyanin biosynthesis, ABA/salt stress/drought, salicylic acid perception, and response to oxidative stress (Table S20). Within these GO categories, we identified the putative tomato ortholog to AtWRKY70 (Solyc03g095770), which functions at the interface of SA and JA signaling (Li *et al*., 2006), in incompatible graft samples at 7 and 14 DAG. Interestingly, the grape ortholog to AtWRKY70 (VIT_08s0058g01390) was previously identified as an upregulated gene in incompatible grape grafts (Assunção *et al*., 2019), making this gene an extremely interesting candidate for future studies in graft incompatibility.

## Discussion

In this study, we use an expanded set of germplasm (four pepper varieties from two different *Capsicum* species) to demonstrate that tomato and pepper are broadly incompatible. This assessment is based on the formation of weak graft junctions and failed vascular reconnections between all tomato/pepper combinations (Fig 1). Notably, previous literature cited the Doux des Landes (DDL) pepper variety as graft-compatible with tomato (Deloire & Hébant, 1982). This historic assessment was likely based on high survival rates and the overall healthy appearance of tomato:DDL combinations. However, we demonstrate that along with all other varieties of pepper tested, tomato-DDL grafts fail to form xylem bridges, and as a consequence, develop biophysically unstable junctions that fail the bend test. Based on our findings, we emphasize the importance of verifying anatomical connectivity when diagnosing graft compatibility, and we recommend additional analyses investigating xylem formation and long-term productivity, to unambiguously assess compatible combinations. For all of the heterografts tested, we also were able to show that vascular bridges were not formed and growth was reduced (Fig. 1-2).

In order to analyze cell death in the graft junction, we tracked the percent of non-viable tissue (NVT) in the junction during the first 3 weeks post-grafting (Fig. 3). All grafts exhibited elevated NVT at 7 days after-grafting (DAG), compatible grafts exhibited significantly reduced NVT at 14 and 21 DAG, while incompatible grafts maintained the same percentage of NVT over time. Programmed Cell Death (PCD) is an important genetic mechanism that allows for selective cell death. In addition to its function in developmental processes (e.g. tracheid and vessel element maturation), PCD plays a central role in specific defense responses, including HR. Due to the high rate of developmental cell death related to vasculogenesis, we were unable to definitively show that PCD caused NVT in incompatible junctions (Fig. 4). To further investigate the cause of sustained NVT in incompatible grafts, we used RNA-seq to analyze the molecular signature of compatible versus incompatible grafts (Fig. 5). In addition to the incompatible grafts displaying a prolonged transcriptional response up to 21 DAG compared to self-grafts, genes upregulated at these later time points were enriched for processes associated with defense responses. Of these upregulated incompatible genes were many NLRs (Fig. 6). NLRs form complexes that monitor both effector presence and effector mediated changes to other proteins; when activated, NLRs trigger ETI and downstream defense responses (Jones *et al*., 2016). NLRs can also self-activate, triggering an inappropriate immune response (Bomblies *et al*., 2007; Tran *et al*., 2017). This phenomenon was originally identified as a type of genetic incompatibility present in F1 offspring of interspecific crosses, leading to the name “hybrid necrosis” (Hollingshead, 1929). Plants executing this immune response display cell death lesions, reduced growth, yellowing, and even complete death (Bomblies & Weigel, 2007). This phenomenon is now attributed to an autoactivated immune response. The neofunctionalization of NLRs in individual species has led to expanded and diverse families, which when crossed can interact deleteriously, activating defense responses in a similar mode to pathogen triggered defense (Tran *et al*., 2017). The expression of NLRs must remain tightly controlled, since upregulation leads to serious growth penalties (Tian *et al*., 2003). Furthermore, overexpression of NLRs can be sufficient to activate autoimmunity (Lai & Eulgem, 2018). Like in other instances of NLR autoactivity, where many NLRs are upregulated, we found significant upregulation of the NLRs from both tomato and pepper (Barragan *et al*., 2021). The *Capsicum* genomes have undergone extensive expansion of the NLR family (Kim *et al*., 2017). It is possible that the phenotypes we observed in incompatible tomato-pepper grafts are caused by NLR-related genetic incompatibility. This would be the first instance of NLR autoactivation being triggered by physical rather than reproductive genomic combinations. Given that the graft junction is composed of an interspecific fusion of tissues, where the genetic information of tomato and pepper are in intimate proximity, it is logical that grafting could elicit a similar response to hybrid incompatibility. Hypersensitive response requires salicylic acid and a core set of genes PAD4, SAG101, and EDS1 in Arabidopsis. We found that most of the orthologs to these HR regulators, in addition to SA responsive genes, were upregulated in incompatible grafts compared to self-grafted controls. Our molecular evidence points to a model in which tomato-pepper graft incompatibility is caused by an immune response activated by incompatible genetic information, which is perceived by differing NLR alleles present in the tomato and pepper genomes.

In addition to the shared NLR-upregulation in tomato and pepper heterografts, we also identified a set of shared orthologs that are upregulated in both the tomato and pepper genomes during incompatible grafting. Further analysis of these shared orthologs may help to identify genetic markers for incompatibility in Solanaceae (Fig. 7).

Next, to explore the genetic fingerprint of graft incompatibility, we compared upregulated genes from compatible grafts, incompatible grafts, and 3 biotic stress datasets (herbivory, fungal infection, and plant parasitism (Fig. 8). We identified an overlap between grafting and these biotic stressors, with a significantly pronounced overlap of upregulated genes between grafting and plant parasitism. Given that the formation of the parasitic haustorium and the graft junction both require inter-specific tissue coordination leading to vascular reconnection, it is logical that the two processes share molecular machinery. The similarity between these two phenomena, both genetically and physiologically, will require future research to fully explore. This analysis also revealed over 1000 uniquely upregulated genes that are expressed in incompatible grafts, including DNA damage repair genes, BRCA1 and BARD1 (Fig 8). Previous work has shown that autoimmune responses caused by NLR overactivation induce DNA damage via EDS1 (Rodriguez *et al*., 2018). Based on these findings we propose that NLR autoactivation is triggered by genetic incompatibility between the diverged immune systems of tomato and pepper, and this immune response triggers a hypersensitive response producing the prolonged accumulation of non-viable tissue in incompatible graft junctions that shares similarities with HR-induced lesions. Further supporting our model, our incompatible grafts shared a unique upregulation of DNA damage repair and HR-related genes that are associated with NLR-mediated autoimmunity. From this analysis, we have identified NLR-mediated genetic incompatibility as a likely cause for tomato-pepper graft incompatibility.

## Supporting Information

Fig. S1 Tomato and pepper heterografts have reduced survival and weak graft junctions.

Fig. S2 Incompatible grafts have reduced secondary growth 30 DAG.

Fig. S3 Tomato and pepper grafts were collected at 7, 14, and 21 DAG for TUNEL assays, trypan blue, and RNA-seq

Fig. S4 Non-viable tissue was quantified in ImageJ

Fig. S5 Heterografted tomato and pepper have consistent growth and persistent non-viable tissue

Fig. S6 All grafted plants have elevated programmed cell death in the graft junction Fig. S7 All grafted plants have elevated programmed cell death in the graft junction (merged)

Fig. S8 Cross-species exudates do not affect callus growth of tomato or pepper Fig. S9 Genetic overlap between tomato and pepper grafts shows scion-stock specificity.

Fig. S10 Scion and stock tissue have distinct upregulated genes at any given time point Fig. S11 Hormonal regulation but not ROS production is upregulated in incompatible grafts

Fig. S12 DNA quality decreases over time in incompatible stocks

Fig. S13 The steroidal glycoalkaloid biosynthesis pathway is significantly upregulated in heterografted grafted scions.

Fig.S14 Biological stressors upregulate distinct and shared genetic responses

Method S1 Plant material and growth conditions Method S2 Grafting

Method S3 Pepper Compatibility Grafts Method S4 Propidium Iodide Staining Method S5 Bend Test

Method S6 Instron three-point bend test

Method S7 DAMP Assay Method S8 Trypan Blue staining Method S9 TUNEL Assay

Method S10 RNA-sequencing and bioinformatic processing Method S11 Orthogroup Parsing

Method S12 Comparative Transcriptomics Method S13 Statistical Analysis

Table S1 Graft survival overtime and manual bend tests

Table S2 Phenotypic data from pepper and tomato compatibility screen Table S3 Instron 3-point bend test results and statistical analysis

Table S4 Non-viable tissue data and statistical analysis

Table S5 Wound exudate and DAMP assay on callus growth and statistical analysis Table S6 *Solanum lycopersicum* (tomato) raw read counts

Table S7 Ca*psicum annuum* (pepper) raw read counts Table S8 Tomato Wald Test output

Table S9 Pepper Wald Test output

Table S10 Genes with upregulation in heterografts based on likelihood ratio test Table S11 GO term enrichment of genes upregulated in heterografted plants as determined by likelihood ratio testing

Table S12 Genes involved in processes of interest which were used to generate heatmaps

Table S13 Alignment rate of RNA-seq libraries

Table S14 Orthogrouping of TAIR10 (Arabidopsis), ITAG4 (tomato), and CM334 (pepper)

Table S15 Shared Orthogroups

Table S16 GO term enrichment from orthogroup overlap Table S17 Genes upregulated following biological stressors Table S18 Genetic overlap and statistical analysis

Table S19 GO enrichment of genes overlap between grafting and biological stressors Table S20 GO enrichment for heterograft-specific upregulated genes

## Acknowledgments

H.R.T. was supported by a United States Department of Agriculture National Institute of Food and Agriculture (USDA-NIFA) Predoctoral Fellowship (2020-67011-31882); M.H.F., A.G., S.P., and M.N. were supported by the National Science Foundation (NSF) (CAREER IOS-1942437). Imaging data was acquired through the Cornell Institute of Biotechnology’s Imaging Facility, with NIH (S10OD018516) funding for the shared Zeiss LSM880 confocal/multiphoton microscope. Access to the Instron Universal testing stand was supported by the Delaware Center for Musculoskeletal Research from the National Institute of Health’s National Institute of General Medical Sciences (P20GM139760). Thank you to Noor AlBader for assistance with data transfer.

## Author contributions

HRT and MHF designed the study. HRT, AG, AH, SY, LE, MN, and ES carried out the experimentation. HRT analyzed the data. HRT wrote the first draft of the manuscript. All authors contributed critically to the final draft and approval publication.

## Data availability

RNA reads collected for this project have been deposited on NCBI GEO (GSE256079). Previously published RNA-seq data used in this research can be accessed on NCBI at PRJNA628162, PRJNA687611, PRJNA756681, PRJNA600385, and PRJNA773605.

